# Transitive reasoning as linear classification

**DOI:** 10.64898/2026.06.24.734346

**Authors:** Vincent P. Ferrera, Samuel Lippl, Kenneth Kay, Fabian Munoz, Yuhao Jin, Greg Jensen, Herbert S. Terrace

## Abstract

Transitive inference (TI) is the ability to reason about transitive relationships in an ordered set of items (e.g., if A>B and B>C, then A>C). TI is widely held to depend on a linear representation of the serial (rank) order of those items. By what computational mechanism is such an ordering constructed during learning, and how is it used to make choices that obey transitivity? Here we take a minimalist approach, applying least-squares estimation (LSE) to a serial learning task commonly used to test TI in humans and animals. In this formulation, LSE computes a linear classifier that maps task conditions onto behavioral outcomes. This algorithm makes no explicit assumptions about transitivity or serial order, yet it reproduces key empirical features of TI; namely, the ability to generalize beyond the training set, and a symbolic distance effect (SDE) in performance accuracy. Applying the classifier to individual items produces an internally ordered representation of rank from which both generalization and the SDE naturally emerge. The approach also yields a decision mechanism, in the form of a differencing operation, for selecting the correct item from any pair. These findings reframe TI as a linear classification problem, challenging conventional assumptions about the cognitive mechanisms required for transitive reasoning.

## Introduction

Computational models of cognitive tasks are often built around complex, neurally inspired algorithms that seek to capture the richness and dynamics of underlying cognitive processes. Such models are validated by their ability to reproduce behavioral and neural responses observed in living subjects. To understand what these models reveal about cognition, however, it is equally important to understand the computational complexity of the task itself. Can simple, ostensibly non-neural algorithms illuminate how cognitive tasks are solved? Here we show that serial learning with transitive inference can be solved by a standard least-squares linear estimation (LSE) algorithm that reduces TI to linear classification. In contrast to neural network and reinforcement learning approaches, which involve many free parameters, the LSE algorithm has no residual degrees of freedom. Yet it exhibits emergent hallmarks of TI performance, including generalization to out-of-training-set comparisons and the oft-reproduced symbolic distance effect in performance accuracy.

Transitive inference (TI) is a classic paradigm widely used to study reasoning in humans and animals. An early example is the problem: “Edith has fairer hair than Suzanne, and Edith has darker hair than Lili. Does Suzanne have darker hair than Lili?” (Piaget, 1928). Such problems can be presented to animals in non-verbal form, and every vertebrate species that has been tested has shown some ability to make transitive inferences (Jensen 2017). The ability to perform TI has been put forth as evidence of logical reasoning in animals (McGonigle and Chalmers 1977).

TI is a component of serial, relational learning and extends to ordered sets much larger than 3 items (Treichler et al., 2003, 2007). A key aspect of TI is that training on a small set of premises about ordinal relationships allows subjects to make inferences about many untrained relations. Thus, if a ranked set of 7 items is written as A>B>C>D>E>F>G, then learning the order of the six adjacent pairs (A>B, B>C, …, F>G) allows one to infer the correct order of the 15 non-adjacent pairs (e.g. B>D). As the set of premises increases linearly, the number of possible inferences grows exponentially.

Over the past few decades, many computational models of TI have been proposed (Wynne 1995; Wu & Levy 2001; De Lillo et al., 2001; Vasconcelos 2008; Kumaran & McClelland 2013; Rombouts et al., 2015; Jensen et al., 2015; Kumaran et al., 2016; Lazareva et al., 2020; Ciranka et al., 2022; Kay et al., 2024; Miconi & Kay 2023; Nelli et al., 2023; DiAntonio et al., 2024, Yang & Maass 2026). Jensen et al. (2019) conducted an extensive analysis of reinforcement learning algorithms applied to TI tasks, finding that the most successful algorithms exploit transitivity to explicitly construct an ordered representation, thereby enabling generalization and symbolic distance effects.

A recent study (Lippl et al., 2024) used mathematical first principles to derive minimal conditions for performing TI and showed that linear stimulus-response mappings necessarily generalize transitively. This analysis raised the possibility of a radically simplified framework for analyzing TI using linear classification. The present study explores that possibility with the aim of clarifying the mathematical basis of transitive inference.

In living subjects, TI is often tested using a limited set of trial conditions, each repeated roughly 10–20 times to obtain reliable estimates of performance metrics such as accuracy and response time. Such repetitive testing is typical of laboratory paradigms with finite sets of stimuli, responses, outcomes, and contexts, and may allow subjects to develop an implicit sense of the overall task structure—including not just the task rules but also systematic relationships between stimuli, responses, and outcomes. TI, like many behavioral paradigms used to probe high-level cognition in animals, can be formalized as a set of stimulus-response-outcome mappings. These mappings can be expressed mathematically as a system of linear equations, admitting a solution—that is, a linear classifier—via least-squares estimation (LSE). The present study applies this framework to a standard TI task and demonstrates that essential features of TI behavior emerge naturally from linear classification. Although TI may appear computationally complex and fundamentally nonlinear, this work establishes that it can be solved by a simple linear mapping and explores some consequences of this observation.

## Methods

### Transitive inference transfer task

In a typical laboratory TI task (**Fig. 1**), a set of 7 images is randomly selected from a large image database. A different set of images is used for each session, and none of these images have been previously presented to the subject. To create an ordered list, the experimenter assigns each stimulus a unique rank, usually indicated by a letter from A to G (**Fig. 1A**). Stimulus rank is never revealed to the subject, nor is there any information about serial order in the spatial or temporal presentation of stimuli.

**Figure 1.**
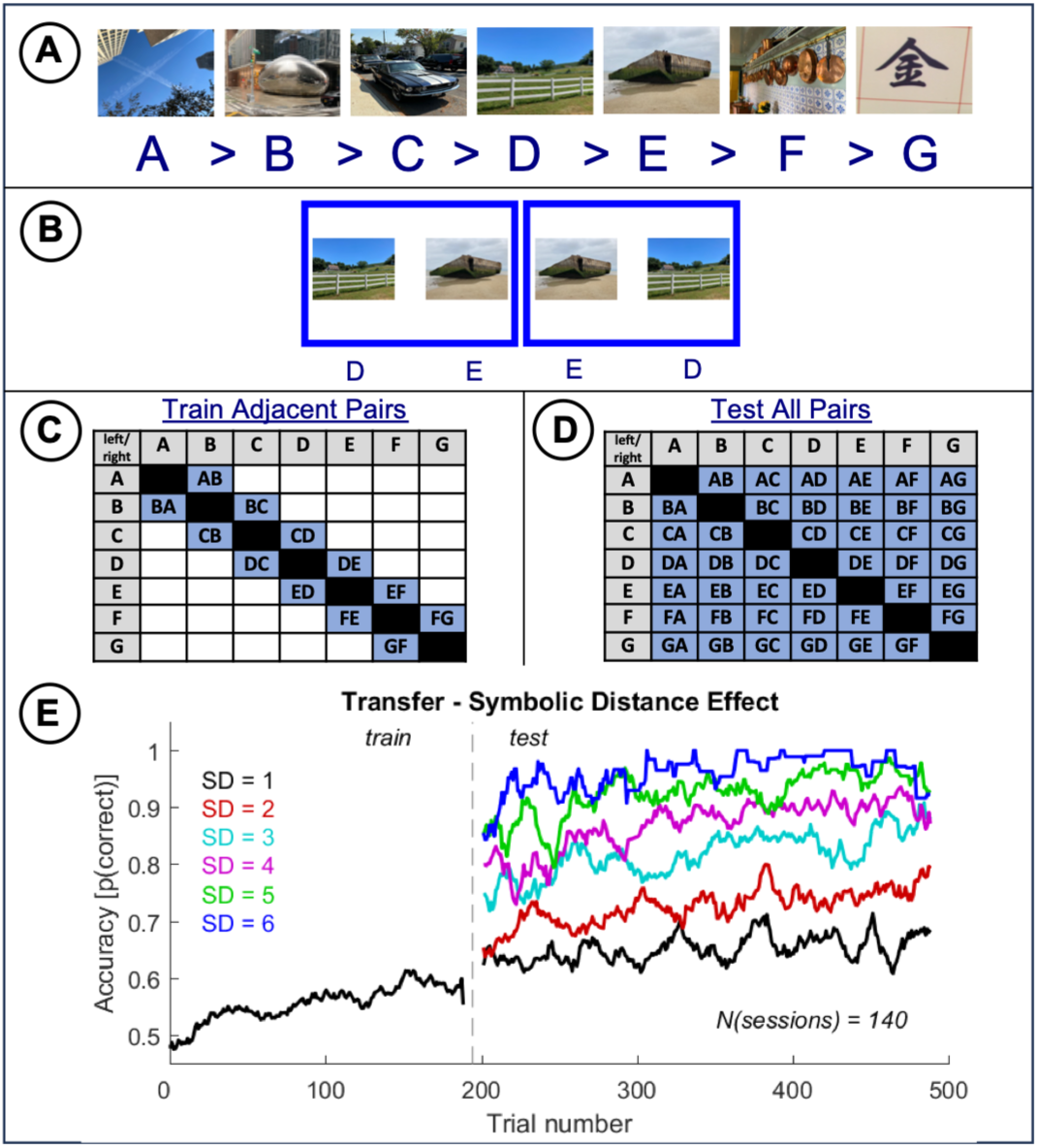
Transitive Inference Transfer Task and Behavior. This task tests subjects’ ability to learn the implied serial order of a set of images from pairwise presentations. **A.** Set of 7 random images is arranged in an ordered list (A>B>C>D>E>F>G). A novel set of images is used for each behavioral session. **B.** A pair of choice trials presenting the pair D/E, with spatial position counterbalanced. **C.** Adjacent pairs used for the training phase of the session. **D.** All pairs used for the testing phase. **E.** Performance accuracy vs. trial number during testing and training. N = 140 sessions.

The task is presented to the subject as a series of trials. During each trial (**Fig. 1B)** two stimuli are displayed equidistant from the center of the screen in opposite locations. The subject’s task is to select the stimulus with the higher rank (lower ordinal) by touching it, moving a mouse cursor, or making an eye movement. Ranks are initially unknown, so initial performance is at chance. Performance improves over the course of a few hundred trials through trial-and-error learning of the correct response to each stimulus pair.

The task is designed to test the transfer of ordinal knowledge from an initial *training* phase to a subsequent *testing* phase. In the training phase, stimuli are presented in pairs (**Fig. 1B**). Each pair comprises stimuli whose ordinal positions are adjacent in the list (A/B, B/C, C/D, etc., **Fig. 1C**). For a 7-item list, there are 6 adjacent stimulus pairs; with positional counterbalancing, this yields 12 unique trials.

In the testing phase of the experiment, all possible pairwise combinations of the 7 stimuli are presented (**Fig. 1D**), one pair per trial. As in the training phase, the correct response is to choose the higher-ranking stimulus. This tests the subjects’ ability to transfer knowledge of list order, acquired during adjacent-pair training, to untrained non-adjacent pairs. For a 7-item list, there are 21 possible pairs (6 adjacent, 15 non-adjacent), yielding 42 unique trials per block with positional counterbalancing. Transfer is defined as correct choice on first presentation of a non-adjacent pair, having been trained only on adjacent pairs. As TI lists grow longer, the number of adjacent pairs increases linearly while the number of non-adjacent pairs grows exponentially; thus, learning a small set of adjacent *premise* pairs allows one to infer a much larger set of ordinal relationships.

Performance of the task by living subjects (3 male nonhuman primates, NHP) is plotted as a function of trial number in **Fig. 1E** (experiments were approved by the Institutional Animal Care and Use Committee of Columbia University. See Munoz et al., 2025 for additional details.) Performance accuracy (percentage of correct responses) was averaged over a total of 140 behavioral sessions from all three animals. There was a gradual improvement in accuracy during the training phase, followed by a dramatic improvement in overall accuracy and a symbolic distance effect at the beginning of testing.

### Simulations

Simulations were run in Matlab 2023b. Code available on request.

## Results

### Intuition for theoretical framework

To develop an intuition for how linear classification can solve TI, we begin with a simple example in which a set of n = 4 stimuli is mapped onto a linear ranking (**Fig. 2**). Each stimulus is characterized by three numbers (***p*** = {*p*_1_*, p*_2_*, p*_3_}), which can be plotted in a 3D space (**Fig. 2A**). These numbers can be thought of as stimulus features. For example, they could represent the color, shape, and size of the stimuli. These properties are assigned at random, and the stimuli are not necessarily ordered in the feature space. However, a vector, ***x***, that maps each stimulus to its rank order, can be found by linear estimation (**Fig. 2B**). This vector assigns a weight to each stimulus feature, ***p***. When the feature vectors are multiplied by the mapping vector (***_p_*** ∗ ***_x_***), the result is a one-dimensional rank ordering of the original stimuli (**Fig. 2C**). The resulting order obeys transitivity even when the original feature-space representation does not. Once ranked, stimuli can be compared across any pair based on relative rank, including non-adjacent pairs not used to generate the ranking. This approach is fully general: given any ranked ordering of the ***n*** items, there exists a mapping vector (classifier) ***x*** that converts the item characteristics, ***p***, into that ranked ordering, provided each item’s characteristics, ***p***, consist of at least *n*-1 values and that each item’s characteristics are distinct (or, more precisely, linearly independent) from those of other items. While **Fig. 2** presents one particular mapping (with the red diamond as the best item and the purple triangle as the worst), other mapping vectors ***x*** could instead be used to obtain any other ranking. The key point is that, given a set of stimuli with measured attributes, a set of weights can always be found that achieves a desired ranking and that set of weights is a linear classifier.

**Figure 2.**
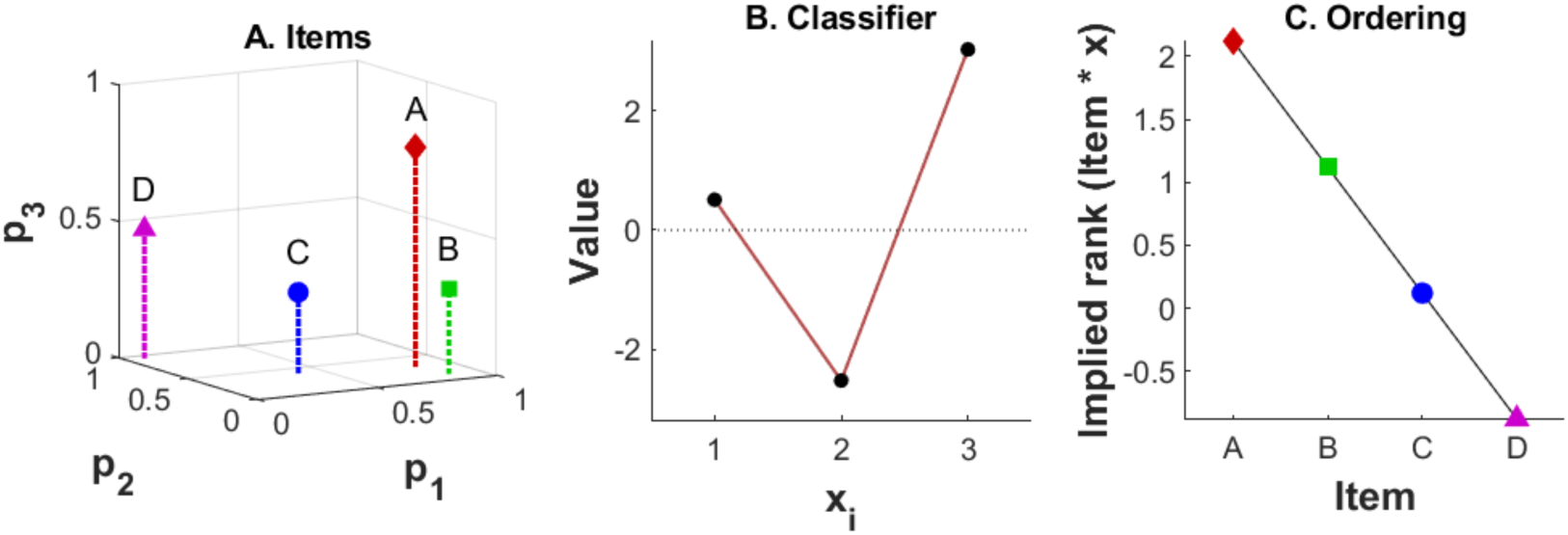
Mapping a set of items with random features onto a linear ranking. **A**) Items with random features plotted in 3D feature space. **B**) Mapping vector (**x**). **C**) Ordering achieved by applying the mapping, **x**, to each item.

### Standard TI Task with Transfer

A typical 7-item transitive inference task can be described as a system of linear equations. To formalize this, a set of *n* = 7 unique stimuli is generated (the analysis generalizes to larger sets). Although laboratory tasks often use pictorial images, a simpler stimulus representation suffices here. Each stimulus is tokenized as a row vector of *p* = 7 elements, where each element is assigned a random value between 0 and 1; *p* is referred to as the *description length*. The stimulus vector can be thought of as a unique “bar code” for each stimulus. There are several possible methods for constructing bar codes from the original stimuli. One is to use a convolutional neural network (CNN) to extract stimulus features, which can then be reduced in dimensionality and orthogonalized by applying singular value decomposition. This is illustrated for a set of 9 pictures in **Fig. 3** using ResNet-50 (He et al., 2016) as the CNN. The orthogonalized output provides a bar code for each of the 9 pictorial stimuli. The set of bar codes, ***A***, is linearly independent (det(***A^T^A***) ≠ 0) and thus satisfies one of the conditions for applying least squares estimation.

**Figure 3.**
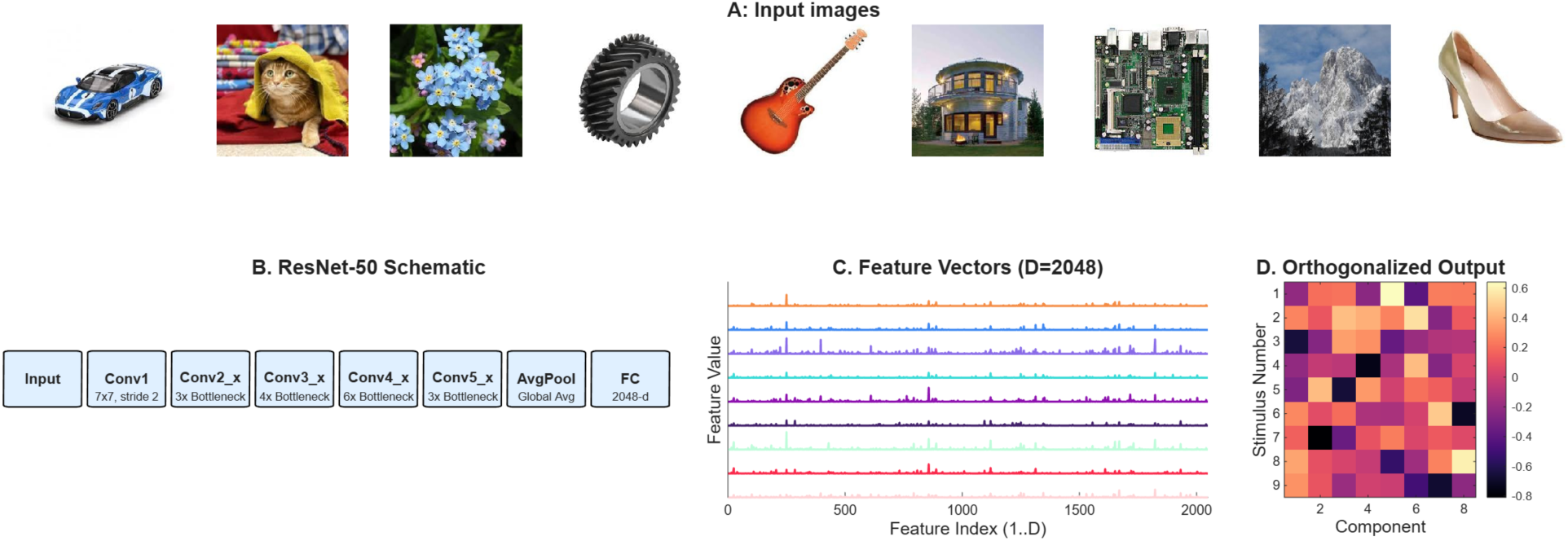
Reduction of input images to stimulus bar codes. **A**) 9 random input images. **B**) Schematic of ResNet-50, a convolutional neural network. **C**) Feature vectors extracted from the original images by ResNet-50. **D)** Stimulus “bar codes” derived from feature vectors by applying singular value decomposition.

Each of the seven stimuli is randomly assigned a unique rank, indicated by the letters A through G, creating an ordered list (**Fig. 1A**). The task is presented to participants as a series of discrete trials. During each trial (**Fig. 1B**), two stimuli are presented. The correct response is to choose the stimulus with the higher rank (e.g., choose A from the pair A/B).

The *training* set comprises the six adjacent stimulus pairs (A/B, B/C, C/D, D/E, E/F, F/G, **Fig. 1C**). Each pair is presented in both spatial configurations—once with the higher-ranked stimulus on the left and once on the right. The *testing* set (**Fig. 1D**) comprises all pairs, adjacent and non-adjacent, presented in both spatial configurations. Spatial counterbalancing ensures the decision is non-trivial: because there are two possible response locations for each stimulus pair, the subject cannot respond correctly without identifying the relative ranks of the presented stimuli.

All of the task elements described above are incorporated into a “task design matrix,” or *embedding*,***W***, that represents the training set as a (2(*_n_* − 1) × 2*_p_*) matrix (**Fig. 4A**). Each matrix row represents a trial condition by concatenating the individual vectors of the two stimuli presented during that trial. The columns of ***W*** are the stimulus features. Thus, the first row is [*A*_1_…*A*_p_, *B*_1_…*B*_p_]. The second row is [*B*_1_…*B*_p_, *A*_1_…*A*_p_]. The third row is [*B*_1_…*B*_p_, *C*_1_…*C*_p_], and so forth.

**Figure 4.**
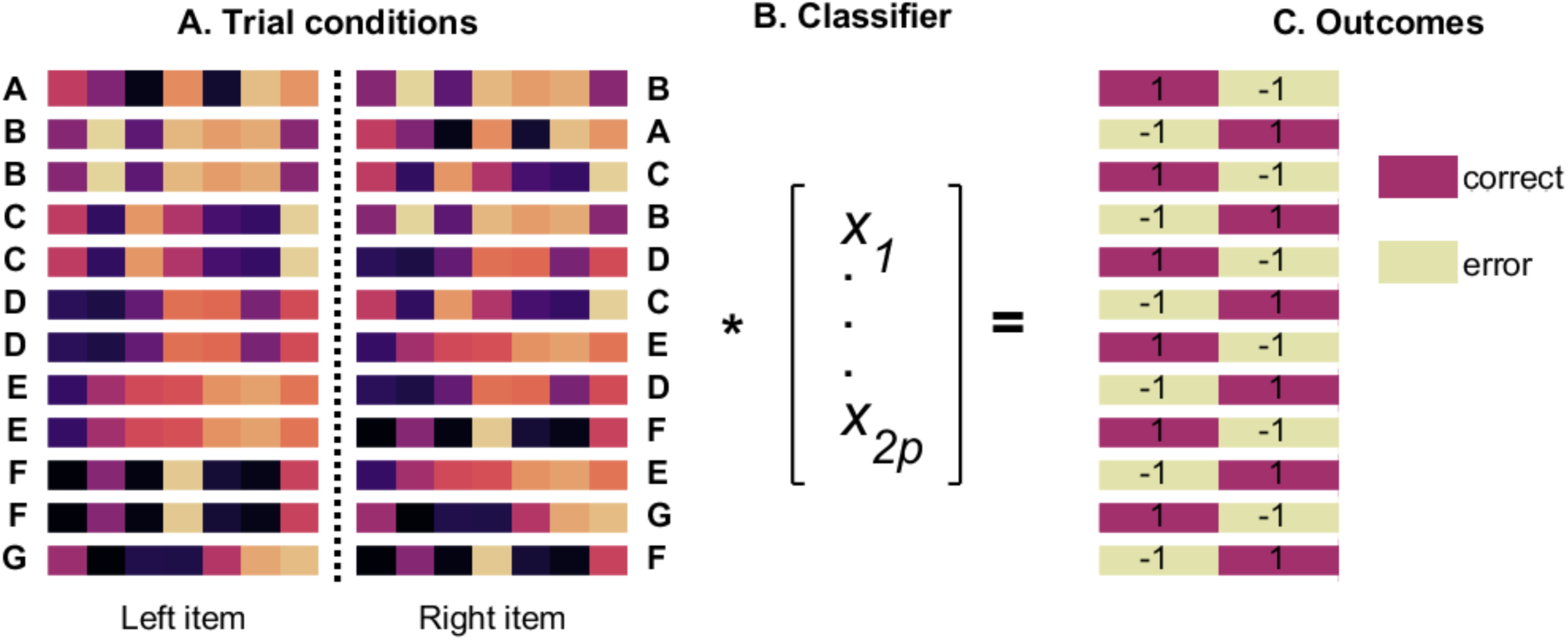
Setup for solving 7-item TI task with LSE. **A**) Task representation (embedding) ***W***. Trial conditions are represented across rows, stimulus features across columns. Each row contains a stimulus pair in a particular spatial configuration. **B**) Mapping vector (***x***). **C**) Outcome matrix (***y***).

The trial outcomes are encoded in a 2*_p x_* 2 matrix, ***y***, where the columns represent the two choices (left and right); 1 and -1 are values that encode the correct and incorrect responses (**Fig. 4C**). This representation assigns a distinct value to each choice option. However, the second column of ***y*** is redundant, being the negation of the first. The outcome representation can thus be simplified to a single column vector where +1 represents correct leftward choices and -1 represents correct rightward choices. This simplification does not affect any of the following results.

The solution to the task is the matrix, ***x***, (**Fig. 4B**) that maps ***W*** onto ***y***. This linear classifier can be found by least-squares estimation (LSE):

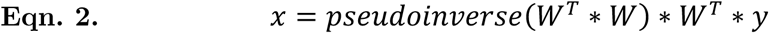

using the Moore-Penrose inverse (Moore 1920, Penrose 1955). The solution, ***x***, is two-fold anti-symmetric; *x*_1…2p,1_ = -*x*_1…2p,2_ and *x*_1…p,i_ = *- x*_p+1…2p,i_, i = 1,2. Hence, we can ignore the second column of ***x*** (*x*_1…2p,2_) and set ***x*** = *x*_1…2p,1_ in the following. The remaining column of ***x*** can be divided into two halves (elements 1 thru *p*, and *p*+1 thru 2*p*) and plotted as in **Fig. 5A**. The solution has *_p_* − (*_n_* − 1) = 0 residual degrees of freedom.

**Figure 5.**
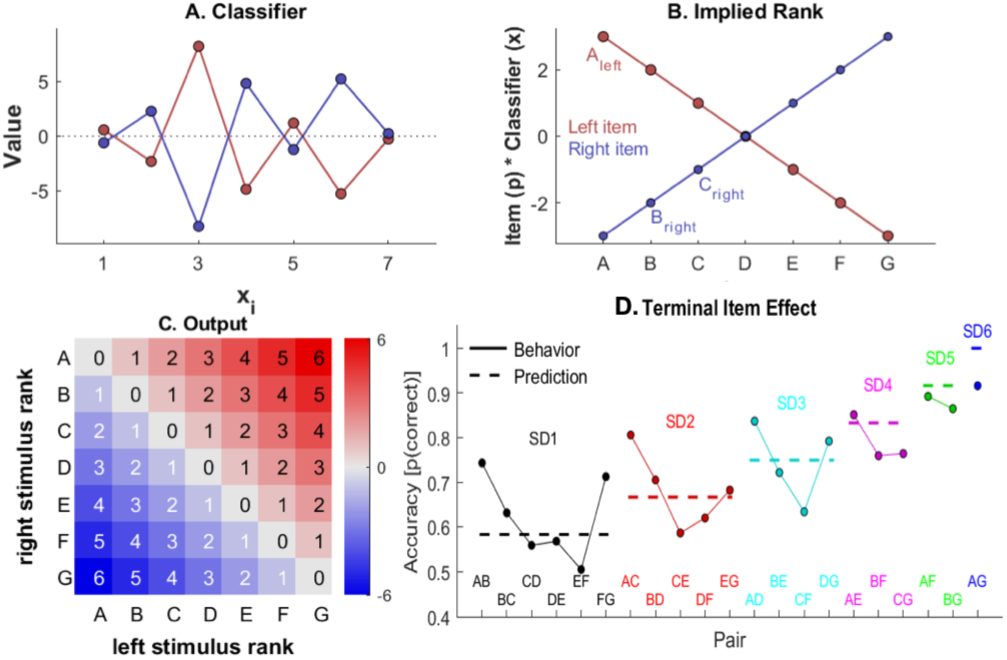
LSE solution to 7-item TI task. **A**. Elements of the mapping vector or “classifier” (***x***). **B**. Implied rank (**Stimulus vector** * ***x***) for left (red) and right (blue) items. **C**. Array of margins for all pairs. Numbers are symbolic distances. **D.** Behavioral accuracy and predicted margin value for each pair in the first block of testing. N = 140 sessions (same data as Fig. 1E)

Performance reaches a plateau when the description length *_p_* = (*_n_* − 1). Increasing the description length further increases the length of the classifier, ***x***, but does not improve or impair performance. Indeed, the bar codes could be replaced with photographic images such as those used in laboratory TI studies, which typically have description lengths in the thousands (proportional to the number of pixels), and performance would remain unchanged.

To understand how the solution to **Eqn. 2** relates to transitive inference, we first note that ***x*** creates an ordered representation of stimulus rank, i.e. the “mental line” that features in nearly every theoretical treatment of TI. To create the mental line, consider the inner product or margin, *M*, of any given stimulus, *U*, with the mapping, ***x***:

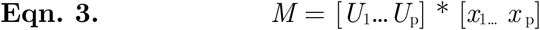

Where [*U*_1_…*U*_p_] is the vector that represents the given stimulus. These inner products decrease in magnitude (**Fig. 5B, red symbols**) with decreasing stimulus rank, demonstrating that weighting the individual stimuli by the linear classifier creates a strictly ordered ranking. This ranking can be used to make choices that obey transitivity.

### Decision Mechanism

The LSE solution incorporates a mechanism for choosing the correct stimulus from each pair. First, note that the classifier, ***x***, is anti-symmetric (**Fig. 5A)** such that the lower half of ***x*** is the negative of the upper half. This points to a differencing mechanism in which the margin (inner product), *M*, for any generic stimulus pair, *U/V*, is equal to:

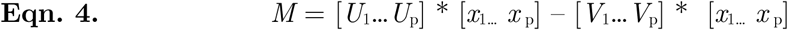

The decision rule is that when the sign of *M* is positive, the stimulus on the left is chosen; when *M* is negative, the stimulus on the right is chosen. Due to the anti-symmetry of the classifier, the subtraction happens automatically by taking the inner product of the trial vector (constructed by concatenating the stimulus vectors *U* and *V*) and the classifier, ***x***. The sign of the inner product determines the choice. This decision mechanism is embedded in the LSE solution and does not require any additional computations.

### Transitive generalization

The linear classifier derived from **Eqn. 2** when the design matrix contains only adjacent pairs generalizes to non-adjacent pairs that were not included in computing the LSE solution.

To demonstrate generalization, the classifier, ***x***, is applied to each non-adjacent pair to produce a scalar margin (*M_i_*_,j_). E.g. for the pair *B/D* with *B* on the left, the margin is the inner product:

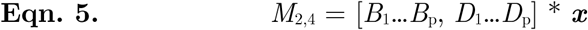

As per the decision mechanism described above, if *M*_i,j_ is positive, then the stimulus on the left is chosen. If *M*_i,j_ is negative, the stimulus on the right is chosen. The complete set of margins for all pairs of stimuli is shown in **Fig. 5C**. The colors indicate that the sign of the margin is correct for each non-adjacent pair (blue = positive = choose the stimulus on the left; red = negative = choose the stimulus on the right). Hence, solving ***_W_*** ∗ ***_x_*** = ***_y_*** for adjacent pairs produces a classifier, ***x***, that generalizes such that taking the inner product of any non-adjacent pair with ***x*** produces a scalar margin whose sign indicates the correct choice.

### Symbolic Distance Effect

The linear classifier produces a symbolic distance effect when applied to all stimulus pairs. Symbolic distance is defined as the difference in ordinal rank between the two stimuli in a pair. The margin for a pair is simply the sum of the margins of the individual items. For example, in **Fig. 5B**, *_Aleft_* + *_Cri_*_gℎ*t*,_ which has a symbolic distance of 2, has a greater margin than *_Aleft_* + *_Bri_*_gℎ*t*,_ which has a symbolic distance of 1. In **Fig. 5C**, the number inside each square is the symbolic distance for the corresponding pair of stimuli. The color saturation of the square represents the margin’s magnitude, bluer for left choices, and redder for right choices. The magnitudes of the margins increase as symbolic distance increases.

When performance was averaged separately for each pair using only the first block of testing trials (42 trials) (**Fig. 5D**), there was a “terminal item effect” (TIE) such that pairs that contained one or both terminal items (A,G) tended to have better accuracy than other pairs with the same symbolic distance. However, the predicted margins did not exhibit a terminal item effect, but were equal for all pairs with the same symbolic distance (**Fig. 5D**.)

### Independence of Trial Order

The LSE solution does not depend on how the rows of the design matrix are ordered. In the example shown above, the rows of the design matrix, ***W***, have a particular order (the first row is always A/B, the second B/A, the third B/C, and so forth). The results do not depend on this specific ordering of rows. The solution depends only on the elements of the stimulus vectors. The rows can be randomly shuffled and LSE will still find a solution that generalizes and shows a symbolic distance effect. In fact, the lines in **Fig. 5A** represent a solution when the design matrix (***W***) was ordered as in **Fig. 2A**, while the filled dots represent the solution when the rows of the same task design matrix were randomly shuffled.

The main results, i.e. generalization from adjacent-only pairs to all pairs, and the symbolic distance effect, do not depend on the outcomes being symmetric (+1 for correct and -1 for incorrect choices). The set of possible outcomes can be any pair of values, e.g. [1 0], as long as the values are distinct.

### Linear separability

The task design matrix, ***W***, can be partitioned row-wise into two trial sets that are linearly separable. Formally, two subsets, **D**₁ and **D**₂, are linearly separable if there exists a weight vector ***x*** and a scalar bias *b* such that:

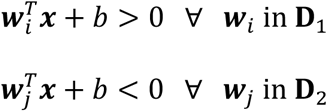

The rows of the design matrix ***W*** used during adjacent-pair training can be divided according to the subject’s choices. Let ***W_+_*** contain the row vectors associated with leftward choices (mapped to positive outcomes), and ***W_-_*** contain the row vectors corresponding to rightward choices (mapped to negative outcomes). Spatial counterbalancing of the task design ensures that these submatrices are perfectly separable through the origin (b = 0), satisfying:

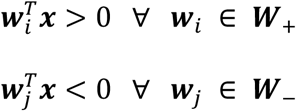

The linear classifier, ***x***, corresponds exactly to the mapping derived in **Equation 2**. This mathematical separability demonstrates that Transitive Inference (TI) tasks can be solved by linear classification architectures, such as single-layer perceptrons and support vector machines, making the problem universally amenable to any linear classifier.

### Dependence on spatial counterbalancing

The design matrix in **Fig. 4A** is spatially counterbalanced and thus encodes complete information about both stimulus location and identity. Spatial information can be reduced by presenting each pair in only one spatial configuration (**Fig. 6A**). To ensure that the decision is non-trivial, the location of the correct stimulus varies randomly (**Fig. 6B**) so that the subject cannot use a default response such as always choosing the stimulus on the right. This approach has been used with monkeys (Jensen et al., 2021) to test their ability to perform TI with minimal training.

**Figure 6.**
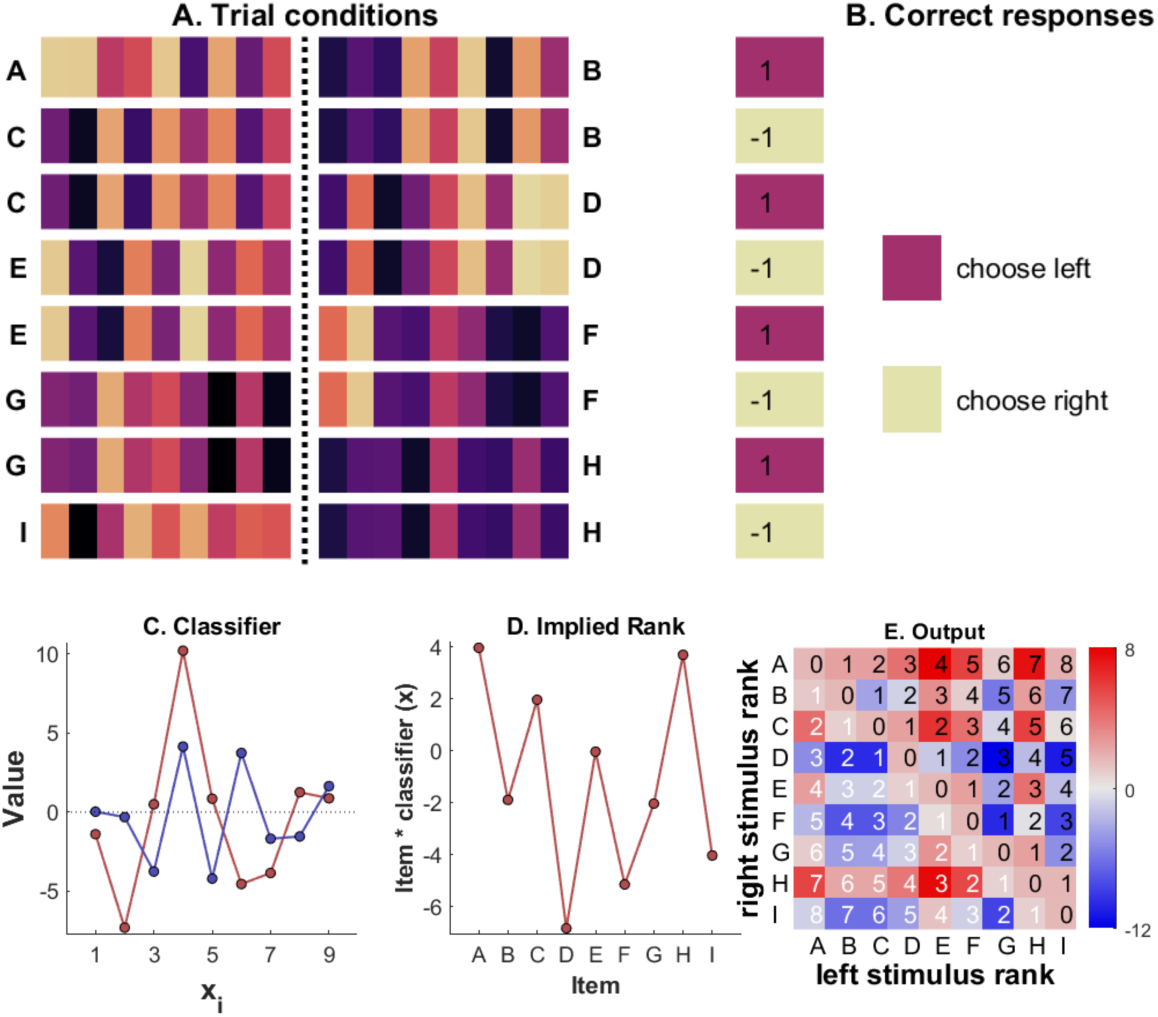
LSE solution to 9-item minimal TI task. **A**) Design matrix. **B)** Outcome vector. **C)** Elements of the mapping vector (**x**). **D**) Implied rank (**Stimulus vector** * **x**). **E**) Array of margins for all pairs.

By removing spatial counterbalancing, the task design becomes indeterminate. There is insufficient information to jointly specify both the relative ranks of the stimuli and the location of the correct response. This would require 2 ∗ (*_n_* − 1) bits of information whereas the outcome matrix, **y**, only provides *_n_* − 1 bits. Consequently, no exact solution exists for this minimal task variant. Nevertheless, LSE can still find solutions (**Fig 6C**), however the implied ranks are not strictly ordered (**Fig. 6D**) and generalization to all pairs is poor (**Fig. 6E**).

### Effects of noise

In living participants, random variability in stimulus processing, response execution, or the stimulus-to-response mapping, due to lapses in attention, memory, or decision-making, is to be expected. The effects of such internal noise can be simulated by adding a random error term, ***e***, to **Eqn. 1**, as follows:

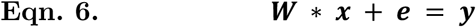

where ***e*** is a matrix comprising random Gaussian deviates with covariance = ***Q***. In this case, the Gauss-Markov solution is:

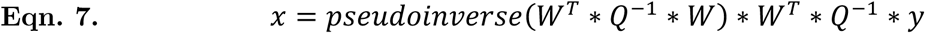

**Fig. 7** presents results from two simulations with noise standard deviation = 0.2, differing only in stimulus vector initialization. In the top row, the classifier, ***x***, (**Fig. 7A**) happens to have approximate mirror symmetry like that shown in **Fig. 5**. This leads to a linear representation of implied rank (**Fig. 7B**), low error (MSE = 0.004, where mean squared error is (***W*** * ***x*** - ***y***)^2^), and appropriate generalization with a symbolic distance effect (**Fig. 7C**).

**Figure 7.**
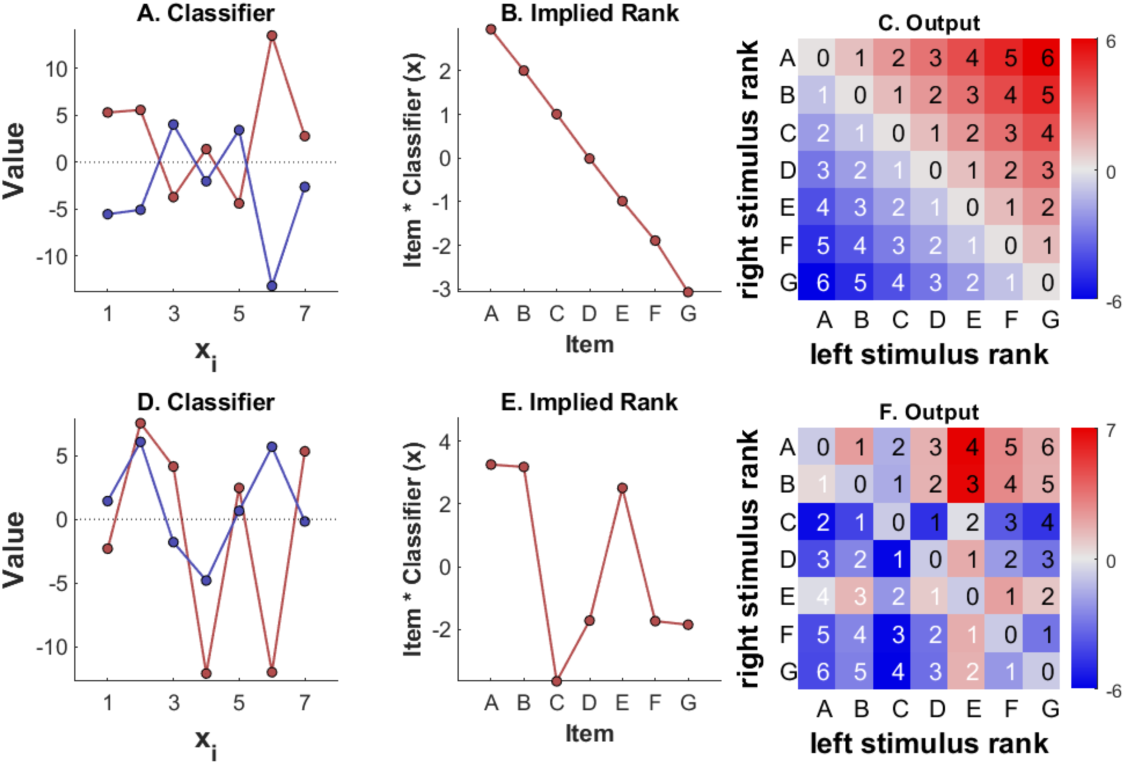
LSE solution to 7-item TI task with added noise. **A,D**) Elements of the mapping vector (**x**). **B,E**) Implied rank (**Stimulus vector** * **x**). **C,F**) Array of margins for all pairs.

In the bottom row, the derived classifier (**Fig. 7D**) lacks symmetry, leading to a large error (MSE = 11.3,), poor reconstruction of implied rank (**Fig. 7E**) and poor performance on all-pairs testing (**Fig. 7F**). These results suggest that the differencing mechanism embodied in **Eqn. 4**, which depends on the mirror symmetry of ***x*,** is critical for successful performance of TI by the LSE algorithm.

### Transverse patterning

Conventional TI has an open-ended topology. The two terminal items have different outcome contingencies compared to the internal list items. For example, internal item C is the correct choice when paired with D, but incorrect when paired with B, yielding a 50% hit rate. By contrast, the first list item (A) is always correct (100% hit rate), and the last item (G) is always incorrect (0% hit rate). This open-loop structure gives rise to a “terminal item effect” (also called an “end-anchor effect”), in which human and animal subjects show markedly higher accuracy on pairs that include a terminal item (Trabasso & Riley 1975).

The loop can be closed simply by adding a comparison between the two terminal items. For a 7-item list, adding the pair G>A transforms the linear topology into a circle—this is called “transverse patterning” (Alvarado & Rudy 1995; Astur & Sutherland 1998). This single addition fundamentally changes the problem: there is no longer a linear classifier capable of mapping the trials onto their outcomes, and symbolic distance becomes ambiguous because it depends on the direction in which the list is traversed.

To confirm the absence of a linear LSE solution for transverse patterns, we constructed a conventional TI list (**Fig. 8A**) and computed the linear classifier using the adjacent pairs (**Fig. 8B**). We show that this classifier creates a ranked representation (**Fig. 8C**), generalizes to non-adjacent pairs, and shows a symbolic distance effect (**Fig. 8D**). Then, without changing the stimulus encoding or any other relationships, we simply added the comparison G>A to close the loop (**Fig. 8E**). This change disrupted performance not only for pairs containing A or G, but for all internal pairs as well, because the classifier (**Fig. 8F**) no longer produced a well-ordered representation (**Fig. 8G**). Consequently, generalization was poor or absent, and no symbolic distance effect was observed (**Fig. 8H**).

**Figure 8.**
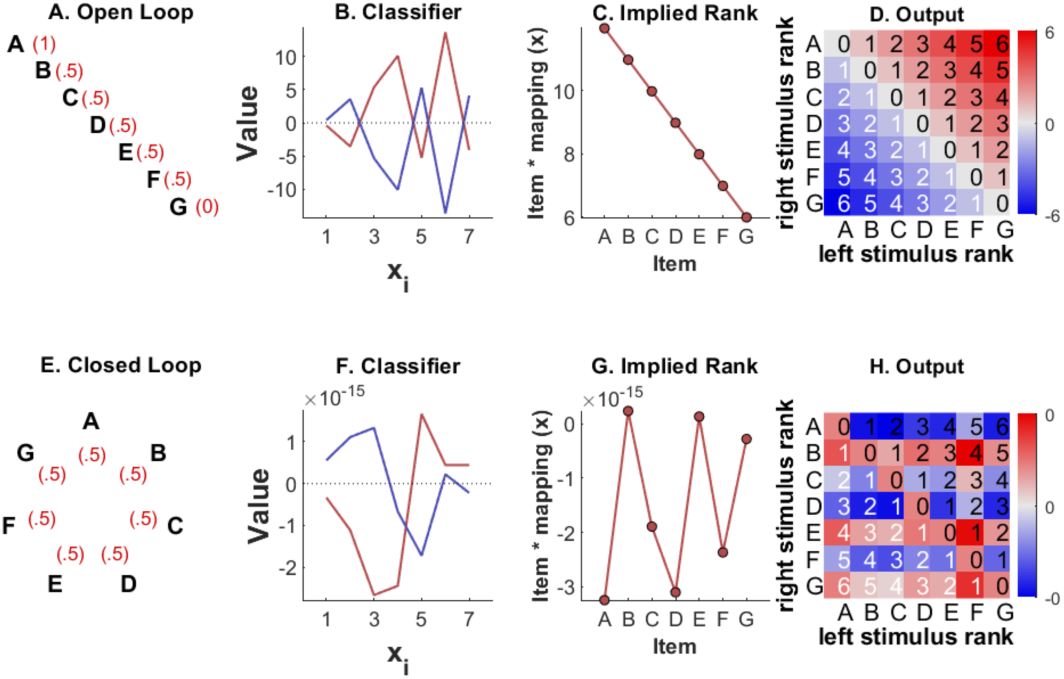
Transitive vs. transverse patterning. **A.** Linear (open) sequence with probability of correct choice for each item (red). **B.** Linear classifer. **C.** Rank derived from linear classifier. **D.** Output for all pairs. Numbers are symbolic distances. **E.** Closed loop formed by adding G>A with probability of correct choice (red). **F.** Classifier. **G.** Rank. **H.** Output for all pairs.

### Perceptron simulation

When provided with the task design and outcome matrices, the LSE solution can be computed in a single step. Living subjects, by contrast, experience one trial at a time and must therefore learn the task incrementally. A well-known approach for solving linear classification problems incrementally is the Perceptron (McCulloch & Pitts 1943; Rosenblatt 1957). In this implementation (**Fig.9A**), each input node (*_xi_*) represented a pair of adjacent items. Input values were initialized randomly. The weighted sum of the inputs was compared to the desired outcome (b), and the resulting error was used to update the weights after each trial. The learning rate was set to 0.05 and the absolute error tolerance was 0.00001.

**Figure 9.**
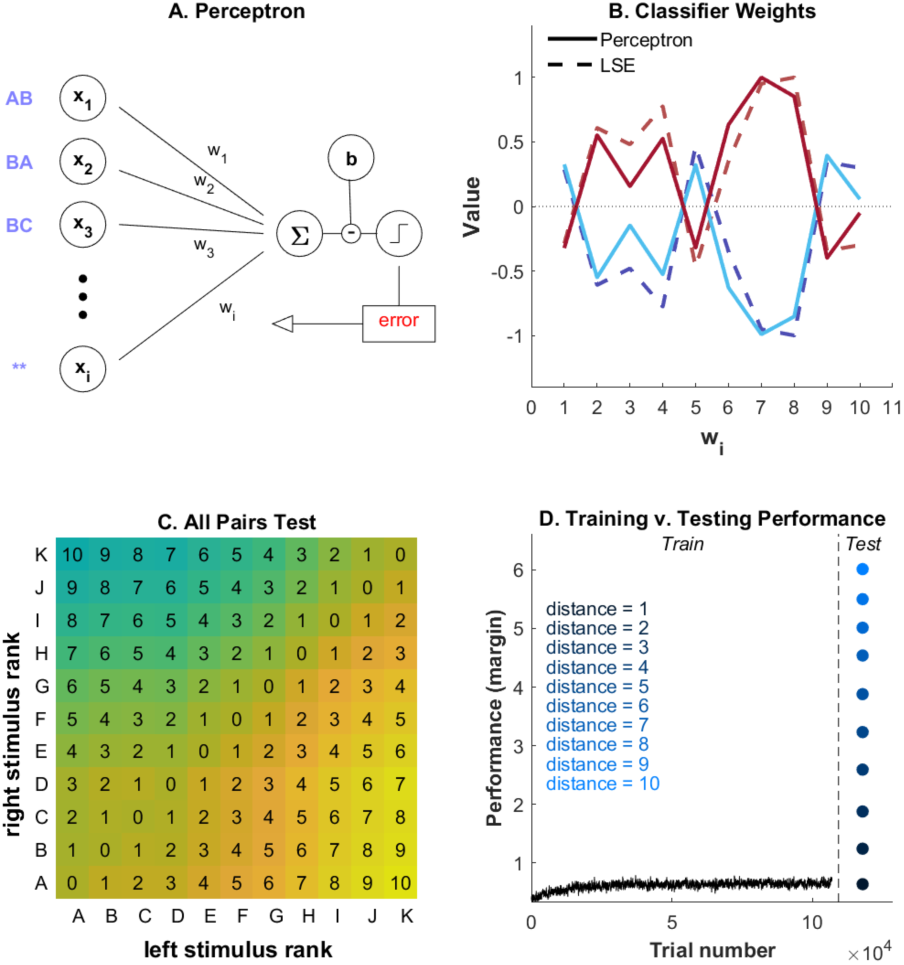
Perceptron implementation. **A.** Perceptron architecture. **B.** Linear classifier weights for Perceptron and Least Squares solution. **C.** Output after training on an 11-item list. **D.** Training and testing performance.

With training on adjacent pairs, the perceptron weights tended to converge to the LSE classifier (**Fig. 9B**). The perceptron solution generalized to all pairs and produced a symbolic distance effect (**Fig. 9C**). During adjacent-pair training, performance, measured as the inner product (margin) between the input pair and the classifier, rapidly reached an asymptote (**Fig. 9D**). Upon transfer to non-adjacent pairs, margins increased markedly and scaled with symbolic distance. This “catastrophic improvement” in performance at transfer is also observed in the behavior of living subjects as shown in **Fig. 1E** and Munoz et al., 2025.

## Discussion

In this study, a transitive inference problem was formulated as a task that could be solved by least-squares linear estimation (LSE), which finds a linear classifier mapping the task representation onto the outcome space. The approach makes no explicit assumptions about transitivity or list order, and requires no iterative training to update weights. Nevertheless, the LSE solution reproduces key features of TI behavior observed in living subjects. (1) A classifier trained on adjacent pairs correctly generalizes to non-adjacent pairs. (2) The magnitude of the decision variable (margin) scales with the symbolic distance between stimuli, yielding a symbolic distance effect. (3) The solution implies a differencing mechanism, also identified in neural network models of TI (Kay et al. 2024). Lippl et al. (2024) formally derive how this mechanism arises from linear stimulus-response mappings. (4) The solution produces a linear representation of rank, i.e. a “mental line,” (Bryant and Trabasso 1971) that is consistent with several computational models of TI.

Whereas Lippl et al. (2024) established analytically that linear stimulus-response mappings necessarily generalize transitively, that result leaves open exactly how such a mapping can be recovered from a standard TI training regime, and whether the resulting classifier reproduces empirical signatures such as generalization, symbolic distance effects, and their failure under transverse patterning that define TI behavior. The present study closes that gap by applying LSE directly to a concrete 7-item transfer task. We show that the minimal linear solution not only exists but is empirically sufficient, requires no free parameters, and predicts exactly where linear classification should fail (transverse patterning). In this sense, the current work operationalizes the Lippl et al. theory rather than re-deriving it.

The way LSE solves TI becomes clear when we examine the inner product of each stimulus with the linear classifier (**Fig. 5B**). The magnitude of these inner products tracks the ranking of the stimuli. This ranking allows the solution to generalize and exhibit a symbolic distance effect. The implied ranking is reminiscent of models that solve TI by implementing transitivity explicitly (e.g., Jensen et al., 2015; Jensen et al., 2019). The explicit application of the transitive rule yields an ordered representation. In the LSE solution, the transitive rule is not explicit; rather, transitivity implicit in the task design constrains the linear classifier so that an ordered representation emerges from solving **Eqn. 1** and then applying the resulting linear classifier to each stimulus.

The approach described here depends on well-known conditions for solving systems of linear equations, i.e. the outcomes must be a linear combination of the task conditions. It is not necessary that the task matrix, ***W***, be square or have a true inverse. As long as the description length of each stimulus is greater than the number of stimuli, the pseudoinverse can find a reasonably good solution and is robust to noise. It is also a condition that the stimulus vectors are not colinear, i.e. the stimulus representations must be distinct and not merely linearly combinations of one another. These constraints place conditions on the neural representation of stimuli that form ordered lists that obey transitivity. It may be the case that TI learning involves the creation of stimulus representations that are linearly classifiable.

The current approach is not intended as a literal model of how the brain performs TI. However, biologically plausible mechanisms for matrix operations such as the pseudoinverse have been proposed (Tapson & van Schaik 2013). More broadly, the LSE approach reveals information that is latent in the task structure and accessible via linear classification and thus amenable to a broad class of models including perceptrons and support vector machines. Transitivity is not assumed *a priori* but emerges from constraints implicit in the task. These results suggest that generalization from adjacent to non-adjacent pairs, and the symbolic distance effect, may be properties of the task structure itself rather than of any particular computational architecture. This view is consistent with recent findings that generic architectures with minimal constraints can perform TI (Kay et al. 2024; Lippl et al. 2024). Subjects who have learned a TI task likely acquire an implicit representation of that structure, and the present results suggest this representation may suffice to account for many serial learning phenomena.

The LSE solution is not incremental: it does not process individual pairs sequentially or update the solution trial by trial; the system of equations is solved simultaneously. However, LSE can be implemented recursively rather than in batch mode. In fact, LSE is effectively implemented by a Perceptron (McCulloch & Pitts 1943; Rosenblatt 1957) with a linear activation function (Beilina 2020). Linear perceptrons can also solve TI (**Fig. 9** and DiAntonio et al., 2024), exhibiting generalization and a symbolic distance effect. As long as the task design is linearly separable, it should be solvable by any linear classification method.

We speculate that the approach described herein can be generalized to any task that can be expressed as a system of linear equations ***W*x*** = ***y***, where ***W*** contains the task conditions and ***y*** represents behavioral outcomes. The utility of this approach is that it distinguishes aspects of behavioral performance that might be expected from the general mathematical structure of the task, such as the symbolic distance effect, from those that may reflect the particular computational mechanisms implemented by a task-solver. The terminal item effect may be an example of the latter.

Cognitive tasks are generally designed to probe specific cognitive functions, but the correspondence between task and function is seldom exact, and validation is necessary. Some cognitive functions are defined entirely by performance on specific tasks; others involve cognitive structures inferred from performance patterns. Computational modeling is an important validation step, but models often presuppose a particular solution strategy. Subjects, by contrast, may be given minimal instruction and are free to adopt any strategy—effective or otherwise. They may arrive at an understanding of the task, or discover strategies, that experimenters never anticipated. For this reason, it is valuable to consider the full range of computational approaches to solving a task, including those that may not initially appear biologically plausible.

## Data availability

Simulation code available on Zenodo

## Competing interests

The authors declare they have no competing interests.

## Acknowledgements

Supported by NIH R01-MH111703 (VPF and HST)

## Author Contributions

The concept for this manuscript was developed by VPF, SL, and KK. The manuscript was written by VPF, SL, KK, FM, YJ, GJ, and HST.

## Notes

### Competing Interest Statement

The authors have declared no competing interest.

### Summary of Updates

This revision has 2 major updates: 1. addition of behavioral data from living (NHP) subjects to illustrate the emergence of the symbolic distance effect at transfer and 2. addition of a CNN to show how real images can be reduced to the "bar code" representation used as input to the least squares estimator.

